# Consistent microorganisms respond during aerobic thaw of Alaskan permafrost soils

**DOI:** 10.1101/2025.08.11.669168

**Authors:** Joy M. O’Brien, Nathan D. Blais, Hannah Holland-Moritz, Katherine L. Shek, Thomas A. Douglas, Robyn A. Barbato, Jessica Gilman Ernakovich

## Abstract

Arctic systems are experiencing warming at four times the rate of the global average, causing permafrost—permanently frozen soil, ice, organic matter, and bedrock—to thaw. Permafrost thaw exposes previously unavailable soil carbon and nutrients to decomposition—a process mediated by microbes—which releases greenhouse gases such as carbon dioxide and methane into the atmosphere. While it is well-established that thaw alters the composition and function of the permafrost microbiome, patterns revealing common responses to thaw across different permafrost soil types have not yet emerged. Here, we address how permafrost thaw impacts microbiome diversity, alters species abundance, and contributes to carbon flux in the Arctic. We sampled peat-like, mineral, and organic-mineral permafrost from three locations in central and northern Alaska and assessed their abiotic soil properties and microbiome characteristics during a 3-month laboratory microcosm incubation. In all sites, prokaryotic biomass increased following thaw, measured as 16S rRNA gene copy number and absolute abundance. This change in biomass was positively correlated with cumulative respiration, indicating an increase in microbial activity post-thaw. We assessed the thaw response of microbial taxa across three sites, identifying taxa that significantly increased in abundance post-thaw. Common responders shared across all sites belonged to the families *Beijerinckiaceae*, *Burkholderiaceae*, *Clostridiaceae*, *Oxalobacteraceae*, *Pseudomonadaceae*, and *Sporichthyaceae*, indicating a common set of taxa that consistently respond to thaw regardless of site-specific conditions. Alpha diversity decreased with thaw across all sites, which likely reflects the increased dominance of specific thaw-responsive taxa, which may be driving post-thaw biogeochemistry and increased respiration. Taken together, we deepen the understanding of different permafrost microbiomes and their response to thaw, which has implications for the permafrost–climate feedback and allows for better predictions of how Arctic ecosystem structure and function respond to change.

## 1 Introduction

High-latitude ecosystems are experiencing rapid environmental change due to increasing global temperatures. These ecosystems are warming approximately four times faster than the rest of the globe (Rantanen et al., 2022) causing degradation of sea ice, glaciers, and permafrost (IPCC, 2019). Permafrost is defined as frozen soil, ice, organic matter, and bedrock that remains at or below 0°C for two or more consecutive years (Van Everdingen, 1998; Schuur et al., 2008). Permafrost harbors a diverse microbiome consisting of dead, dormant, and active microbes across bacterial, archaeal, and fungal lineages (Burkert et al., 2019; Hansen et al., 2007; Steven et al., 2006; Waldrop et al., 2025). Permafrost covers approximately 15% of the northern hemisphere (Obu, 2021) and contains approximately twice as much carbon (C) as there is in the atmosphere (Hugelius et al., 2014; Nadeem et al., 2024), making it a critical component of the global climate system. For some microbes, permafrost soil carbon (C) and nitrogen (N) are largely inaccessible and not easily metabolized due to the frozen conditions, however when permafrost thaws, soil temperature and moisture, as well as the accessibility of C and soil nutrients increase (Ernakovich et al., 2017; Schädel et al., 2016). The release of C following thaw as particulate organic matter (POC), dissolved organic carbon (DOC), and/or mineral-associated organic matter (MAOM) can increase microbial activity and decomposition leading to increased microbial respiration and biomass (Clein & Schimel, 1995; Elberling & Brandt, 2003; Ernakovich et al., 2017; Mackelprang et al., 2011; Schädel et al., 2016). As microbes break down organic matter in thawing permafrost, they produce carbon dioxide, methane, and nitrous oxide as byproducts of aerobic and anaerobic respiration (Ernakovich et al., 2017; Hopkins et al., 2013). These greenhouse gases are released into the atmosphere and contribute to a positive carbon–climate feedback further accelerating global warming (Schädel et al., 2016; Schuur et al., 2015). Therefore, understanding how changes in microbial abundance, composition, and respiration occur is imperative to accurately predict the contributions of permafrost thaw to atmospheric carbon dioxide and further predictions of climate change in the Arctic (Graham et al., 2012).

Previous permafrost thaw microcosm experiments (e.g., controlled temperature incubations) have shown significant changes in the factors affecting the composition of the permafrost microbiome from before to after thaw in a relatively short time period (weeks to months), (Barbato et al., 2022; Doherty et al., 2020; Ernakovich et al., 2017; Mackelprang et al., 2011; Ricketts et al., 2020). For example, in an incubation of soils from a peatland-dominated permafrost site in northern Sweden, depth and temperature were significant drivers of community composition post-thaw, which was also strongly influenced by drift (Doherty et al., 2020). In contrast, corresponding active layer soils were dominated by deterministic assembly, in which nonrandom selection by abiotic and biotic conditions drives composition, which implies a predictable post-thaw community composition (Doherty et al., 2020; Ernakovich et al., 2022). Understanding the consistency and predictability of thaw response is important because it allows for better predictions of the functional capabilities of the post-thaw community.

Knowing which microorganisms proliferate and increase in abundance following thaw could help predict the post-thaw permafrost microbiome composition and function. Microbes that increase in abundance following thaw can be classified as “thaw responders” — a distinct set of taxa that respond similarly to permafrost thaw. Previous research has shown that permafrost microbes exhibit variations in temperature tolerance, and there is potential for the pre-frozen microbiome to respond favorably when conditions improve (Ernakovich et al., 2017, 2022; Waldrop et al., 2023). For instance, one permafrost incubation study identified the emergence of nitrogen-fixing methanogens post-thaw along with an increase in members of the phyla Actinobacteria, Proteobacteria, Bacteroidetes, and Firmicutes (Mackelprang et al., 2011). Another study observed a positive correlation between Alphaproteobacteria and the quantity of mineralized carbon, along with heightened microbial activity in response to readily available nutrients. This study also reported an increase in the abundance of Betaproteobacteria, Gammaproteobacteria, and Bacteroidetes (Ricketts et al., 2020). However, whether there are a common set of microbes or functional groups that increase consistently with permafrost thaw is unknown. Therefore, seeking out the identity of “thaw responders” has the promise to enable understanding of not only post-thaw community composition, but also post-thaw gas fluxes. While permafrost encompasses a diverse range of soils, from mineral to organic, which support varied microbiomes, (Ping et al., 2015; Waldrop et al., 2023, 2025); looking for common responders is a promising way of advancing knowledge across this diversity.

The goal of this study was to understand how permafrost thaw impacts the microbiome and its contribution to carbon dioxide flux, using microcosm experiments with samples from three sites in Alaska. We performed a 96-day thaw microcosm experiment with permafrost from three different sites (ranging from mineral to organic permafrost and varying in carbon content). Microbial respiration was measured throughout the incubation. We determined bacterial abundance via quantitative polymerase chain reaction (qPCR) and community composition via amplicon sequencing of the 16s rRNA gene before and after thaw. We hypothesized that: 1) bacterial abundance and respiration would increase following thaw across sites and 2) a core group of thaw responders would consistently emerge and increase in absolute abundance across sites post-thaw.

## 2 Materials and Methods

### 2.1 Site descriptions, sample collection, and processing

Permafrost samples were obtained from three locations in Alaska and stored at several institutions prior to being shipped to the University of New Hampshire (Table 1). Two permafrost cores were collected per site. One site is located above the U.S. Army’s Cold Regions Research and Engineering (CRREL) Permafrost Tunnel 10 km north of Fairbanks, AK. It represents syngenetic, high carbon and high ice content predominantly eolian permafrost with a typical active layer of ∼60 cm (Douglas et al., 2021) and is located along a gently sloping upland area. Cores from above the tunnel were collected on October 4, 2020. The second site is the CRREL Farmers Loop Experimental Station in Fairbanks, AK which represents a lowland syngenetic ice rich permafrost that has a higher sand content than the Tunnel. Active layer depths are typically ∼80 cm at this site (Douglas et al., 2020). Cores from Farmers Loop were collected on May 26, 2018. The two Fairbanks area sites represent discontinuous permafrost. The third location is from the Barrow Experimental Observatory (BEO), located in an extremely low gradient plane 5 km southeast of Utqiaġvik, AK. Permafrost at the BEO is continuous (up to 400 m thick) and is comprised of sand, silt, and peat layers with typical active layers of ∼ 45-50 cm and 50.8 cm at the time of collection (Douglas & Blum, 2019; Nelson et al., 2008; Thurston et al., 2025). Cores from BEO were collected on September 26 and 28, 2018. All three sites contain ice rich permafrost that can be broadly described as pore-ice cemented soil particles with massive ice wedges. This type of permafrost is carbon rich and is susceptible to the formation of large thermokarst features upon thaw due to the high ice content.

**Table 1.** Permafrost sample collection metadata. Mineral permafrost indicates an organic layer < 40 cm. Organic permafrost indicates organic layer > 40 cm.

| Sample site in AK | Date of sampling | Coordinates | Description | Permafrost donor, Storage location prior to this study |
| --- | --- | --- | --- | --- |
| Above CRREL Tunnel | October 4, 2020 | 64.951147N, 147.620516W | Mineral permafrost (% C < 20) | Tom Douglas, CRREL |
| Farmers Loop (FL) | May 26, 2018 | 64.874775 N, 147.681356 W | Organic/peat-like permafrost (%C > 20) | Taylor Sullivan, Andy Parsekian, University of Wyoming |
| Barrow Experimental Observatory (Utqiagvik) | September 24, 26 2018 | 71.323 N, 156.6114 W | Organic and mineral mix/sandy and silty cryoturbate (%C > 20) | Robyn Barbato, CRREL |

All permafrost cores were collected using a gas-powered 8 cm diameter SIPRE corer (Jon’s Machine Shop, Fairbanks, AK). Cores were approximately 1 meter in length (typically in four ∼25 cm core sections). To protect permafrost samples from contamination, all personnel wore coveralls (Tyvek, Wilmington, DE) and nitrile gloves during sample collection. At all three sites the seasonally thawed active layer soil was removed using shovels and then cores were collected. Samples were kept frozen in the field and remained frozen during transport from Alaska to various institutions to the University of New Hampshire (UNH; Durham, NH). When not in transport samples were stored at −20°C. At UNH, permafrost samples were aseptically prepared for incubation in a freezer room (7°C). Sterile conditions were maintained by cleaning the room, surfaces, and tools with 70% ethanol and 10% bleach. Personnel wore coveralls and nitrile gloves. To remove potential contamination, a sharp stainless steel paint scraper sterilized with 70% ethanol was used to scrape off the outer 0.5 cm of each core. After scraping for sterilization, we sectioned off each core with a sterile PVC wire saw for homogenization. Scraped and sectioned cores, approximately 25 cm in length, were placed in Whirl-Pak bags and stored at −20°C until further use.

Each sample was removed from its Whirl-Pak bags placed into sterile (autoclaved) canvas bags (approx. 35.5 cm long x 12.7 cm wide) and hit with a hammer (sterilized) until fully homogenized. Canvas bags were laundered and autoclaved between use of each sample. Four replicate subsamples (20-40 g) were sampled from each homogenized core for a total of 24 jars for incubation (3 sites x 2 cores per site x 4 subsamples per core = 24 samples). Additional sub-samples were retained and stored at −20°C as a pre-thaw comparison. Due to limited amounts soil, we prioritized replication in pre-thaw microbial analyses over soil analyses saving one pre-thaw subsample per core for soil analyses (see below) and four pre-thaw sub-samples per core for microbial community analyses.

### 2.2 Permafrost incubation

The subsamples were incubated at 2°C for 96 days to gently induce thaw (Figure 1). We chose an incubation temperature of 2°C to mimic near-freezing temperatures typical of early thaw in Arctic regions. Each subsample was aseptically placed into a specimen cup and then in a mason jar fitted with a luer lock to allow for gas measurements throughout the incubation. To avoid water saturation, three holes and glass fiber filter paper were placed in the bottom of the specimen cup which was then placed on an inverted tin foil weigh boat inside the mason jar. This was done to ensure that the incubation remained aerobic. Gas measurements were collected on days 0, 1, 2, 4, 6, 9, 12, 14, 19, 26, 30, 36, 44, 51, 58, 65, 74, 83, 92, and 96 and analyzed for microbial respiration on a Picarro G2201-I cavity ring down spectrometer (Picarro, Santa Clara, CA, United States). All materials involved in the incubation were properly sterilized prior to set-up. In order to ensure that the CO_2_ concentration was below 2% and that the jars remained aerobic, the jar headspace was flushed with Ultra Zero air (Airgas, Dover, NH) for 10 minutes after gas measurements. Jars were randomly arranged in the incubator after each measurement to reduce any unintended treatments. Respiration rate was calculated as µg CO_2_-C g^−1^ dry soil h^−1^. Dry soil weight was calculated using the gravimetric water content of the samples. Cumulative respiration was calculated as µg C-CO_2_ g^−1^ dry soil.

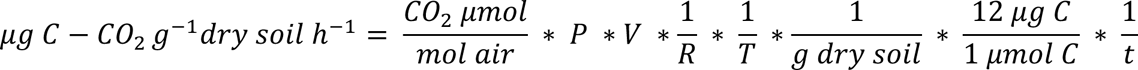

**Figure 1.**
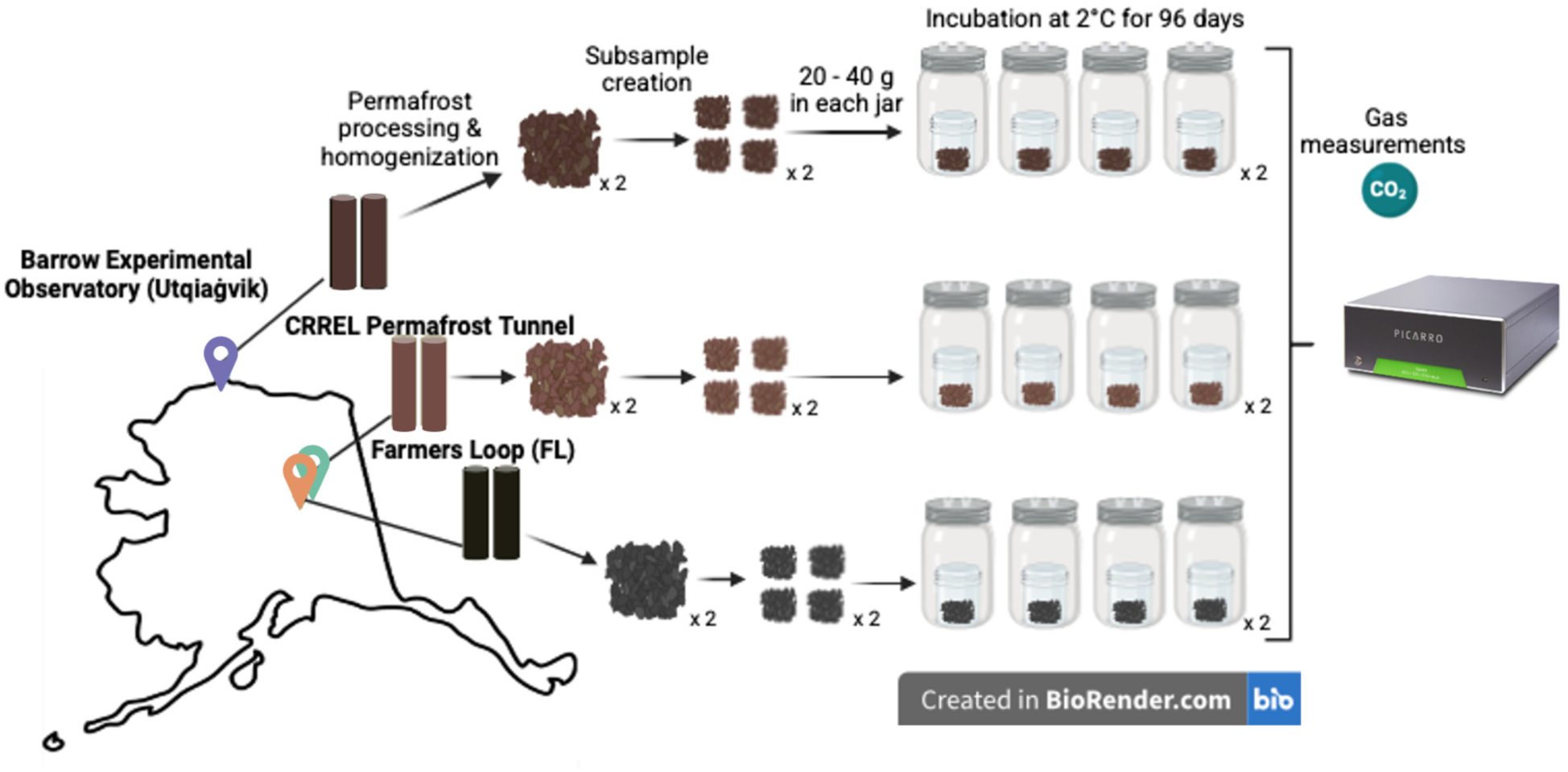
Permafrost incubation experimental design. Two permafrost cores were collected from three different sites in Alaska, USA. Each core was processed, homogenized, placed into jars, and incubated at 2°C for 96 days with intermittent gas measurements (x 2 notation indicates that the process was performed for each core).

We calculated respiration rate where *P* is atmospheric pressure in atm, *V* is the volume of headspace in L, *R* is the ideal gas constant in L atm K_-1_ mol_-1_, *T* is incubation temperature in K, and *t* is incubation length in hours from most recent flush.

### 2.3 Soil analyses

After 96 days of incubation, the thawed permafrost subsamples were destructively sampled for downstream analyses. Pre-thaw soil and post-thaw soil was weighed for the following analyses: 0.25 g for DNA extraction, 10 g for gravimetric water content (GWC), pH, and electrical conductivity (EC). To measure GWC, 10 g of soil was placed in a drying oven at 105°C for 48 hours and weighed after the 48-hour drying period (Gardner, 1965). Soil was then transferred to a 50 mL falcon tube for pH/EC analyses (Rayment & Higginson, 1992). For both the pH and EC analyses, 25 mL of Milli-Q water was dispensed into to each tube and the tubes were placed on a tabletop shaker for 1 hour at 115 rpm. EC was measured first on an Accumet AB30 Conductivity Meter with an Accumet glass body conductivity cell 0.1 cm. pH was measured on an Accumet basic 15 pH meter (Thermo Fisher Scientific, Waltham, MA, United States). Samples that were analyzed for combustible total carbon and nitrogen were measured on dry soil using combustion analysis in replicates of three Costech ESC 4010, Valencia, CA, USA). Combustion occurred at 950°C. Acetanilide was used for calibration in addition to the blanks. Instrument check standard occurred every 12 samples. R2 value for N (whole range) was 0.999651 and C (whole range) 0.999848.

### 2.4 Microbial community analyses

DNA was extracted from the pre-thaw and post-thaw soil permafrost subsamples using the Qiagen PowerSoil Pro Kit (Qiagen, Hilden, Germany) following the manufacturer’s protocol. DNA was then amplified in 12 µL reactions using PCR with the following primers and conditions: 515F with Nextera adaptors (5’TCGTCGGCAGCGTCAGATGTGTATAAGAGACAGGTGYCAGCMGCCGCGGTAA) and 926R with Nextera adaptors (5’GTCTCGTGGGCTCGGAGATGTGTATAAGAGACAGCCGYCAATTYMTTTRAGTTT) of the V4 regions of the 16S rRNA gene to identify bacterial and archaeal communities (Parada et al., 2016; Quince et al., 2011). The PCR reactions were set up as follows: 6 µL DreamTaq Hot Start Green (Thermo Fisher Scientific, Waltham, MA, United States), 2.6 µL of a sterile water solution containing 0.5 µg/µL Bovine serum albumin (BSA) to reduce potential PCR inhibitors (Kreader, 1996), 0.7 µL forward primer (5 µM), 0.7 µL reverse primer (5 µM), and 2 µL template DNA, for a 12 µL reaction total. The thermocycler conditions for 16S rRNA amplification were as follows: enzyme activation at 94°C for 3 min, 35 cycles of denaturation at 94°C for 45 s, annealing at 50°C for 60 s, and extension at 72°C for 90 s, and lastly a final extension at 72°C for 10 min. PCR products were then visualized via gel electrophoresis. During each round of DNA extraction, we included two empty tubes to serve as blanks. All samples amplified successfully and were sent to the University of New Hampshire Hubbard Genome Center, Durham, NH for multiplexing and sequencing on an Illumina NovaSeq 6000 (Illumina, San Diego, CA, USA) with 2×250bp chemistry. A portion of the samples were resequenced to achieve acceptable sequencing depth (> 20,000 reads per sample). To control for run-to-run biases, several samples that were sequenced to sufficient depth on the first run were also resequenced on the second run.

Quantitative PCR (qPCR) was performed on all previously extracted DNA using a Bio-Rad CFX96 thermocycler (BioRad, Hercules, CA, USA). We aimed to quantify the 16S rRNA gene of the V4 region using the aforementioned primers (515F and 926R). DNA samples were normalized to 10 ng of DNA per 46 µL of molecular grade water. Reaction volumes of 15 µL were set up as follows: 7.5 µL SsoAdvanced™ Universal SYBR® Green Supermix (BioRad, Hercules, CA, USA), 1.125 µL (10 µM) forward primer, 1.125 µL (10 µM) reverse primer, 3 µL template DNA, and 2.25 µL molecular grade water. The thermocycling protocol was carried out as follows: polymerase activation at 95℃ for 4 min, denaturation (40 cycles) at 95℃ for 30 s, annealing at 53℃ for 30 s, and extension at 72℃ for 60 s. Standard curves were composed of five 10-fold serial dilutions (0.836 to 0.00000836 ng/µL) of genomic DNA from *Escherichia coli* k-12 (Blattner et al., 1997) that was extracted from pure culture (Carini et al., 2016). All samples and standards were run in triplicate. All reaction efficiencies were >104% and all R^2^ were >0.986. Results were standardized to the original mass of soil and are reported as gene copies g^−1^ soil.

### 2.5 Sequencing and statistical analyses

The two sequencing runs were independently run through the DADA2 pipeline for denoising and assembly into amplicon sequence variants (ASVs) with modifications to the error rate learning step for NovaSeq quality scores (Callahan et al., 2016; Teixeira et al., 2024). Briefly, primer sequences were removed with cutadapt, and reads were filtered and trimmed for quality (quality plots determined the number of base pairs to trim). The resulting ASV tables were merged using the ‘mergeSequenceTables’ function within DADA2. Chimeras were removed and taxonomy was assigned with Silva db v138. To account for sequence run-based differences in base calling, ASVs in the merged sequence table were aligned and then clustered at 99% sequence similarity using the ‘DECIPHER’ and ‘speedyseq’ R packages (Wright, 2016; McLaren, 2020). We reassigned taxonomy on the clustered ASVs using the ‘assignTaxonomy’ function in DADA2 (Callahan et al., 2016). The merged sequence table contained 19,617 ASVs. The final ASV table after clustering at 99% similarity contained 17,073 ASVs across 52 samples (after the second merge of duplicate samples) and was used for all further statistical analyses in the ‘phyloseq’ (McMurdie & Holmes, 2013) and ‘vegan’ packages (Oksanen et al., 2022). All analyses were performed in R (version 4.2.0; R Core Team, 2021). Taxa unassigned at the phylum level were removed (1025 removed) in addition to reads that were identified as mitochondria and chloroplasts (6835 which was approximately 42.6% of the total reads). DNA and PCR extraction blanks were removed from the sequencing data set after determining that there was no contamination (5 sample blanks removed from the dataset). We rarified the samples to a depth of 5000 (this depth ensured that majority of the samples were retained). We removed samples that experienced lab error (FL1C4 REP 4 pre- and post-thaw samples).

Following sequencing, we employed a number of statistical analyses to evaluate the collected data. One-way analysis of variance (ANOVA; in the ‘vegan’ package) was used to test for differences in the post-thaw soil abiotic factors by site such as pH, EC, total carbon, and total nitrogen, followed by post-hoc analysis of between group differences using Tukey HSD tests (in the ‘stats’ and ‘multcompView’ packages). For abiotic data that failed the assumptions of normality (Shapiro-Wilk test in the ‘stats’ package) and homogeneity of variance (Levene’s test), we used the non-parametric Kruskal-Wallis test (in the ‘stats’ package) for significance followed by a Dunn test (in the ‘FSA’ package) with p-values adjusted with the Holm’s method for post-hoc comparisons. Correlations to relate gene copy number to cumulative respiration were performed using the Spearman method via the “cor.test” function (in the ‘stats’ package). Boxplots and thaw response dot plots were made in ggplot2 (Wickham, 2016). Diversity metrics were calculated using the Shannon and Simpson indices using the ‘vegan’ package followed by further analysis with the Kruskal-Wallis and Dunn tests. We measured community dissimilarity between prokaryotic communities using Bray-Curtis dissimilarity measure and visualized using principal coordinates analysis (PCoA) ordination plot. Differences in community dispersion were assessed using the ‘betadisper’ function (homogeneity of dispersion) followed by a permutation test of multivariate homogeneity of groups using the ‘permutest’ function (n = 999) and Tukey’s HSD test to determine pairwise comparisons across site. PERMANOVA was performed using the ‘adonis2’ function in the ‘vegan’ package to test for differences in microbial communities by site and pre-post-thaw treatments. The relative abundance of taxa were scaled accordingly by the 16S rRNA gene counts from qPCR to obtain absolute microbial abundance (Jian et al., 2020). Response ratio analysis (Hedges et al., 1999) was used to identify the absolute abundance of ASVs that increase due to permafrost thaw (Fan et al., 2022). The ASV data frame used in the response ratio analysis was scaled up by 0.01 to eliminate zeros while maintaining the presence-absence relationship. Log response ratios (RR) were calculated as the natural log of post-thaw abundance divided by the pre-thaw abundance. Standard errors for each RR were estimated followed by calculating Z-scores and a two-tailed test. P-values were corrected for false positives in multiple comparisons using the Benjamini & Hochberg false discovery method (Benjamini and Hochberg, 2000). Individual ASVs were considered significantly responsive if FDR-adjusted p < 0.05. We assessed response ratios that showed an increase of two-fold or more post-thaw.

## 3 Results

### 3.1 Pre-thaw and post-thaw abiotic analyses

Soil abiotic properties varied from pre-thaw to post-thaw and within sites (Table 2). Pre-thaw pH across all sites was acidic and ranged from 4.16 to 4.91, and post-thaw pH ranged from 3.94 to 4.88 across all sites. Post-thaw pH significantly decreased between Utqiaġvik and CRREL (TukeyHSD p-value = 0.001) as well as Utqiaġvik and Farmer’s Loop (TukeyHSD p-value = 0.001). Electrical conductivity (EC) ranged from 206.3 to 441.6 (mS/cm) before thaw. Following thaw, EC (228.7 to 478.5 mS/cm) was significantly less at CRREL compared to Utqiaġvik (ANOVA F = 13.95, p-value =0.0303, TukeyHSD p-value =0.02). Pre-thaw total combustible C (% C) differed across sites, ranging from 1.8% in CRREL to 39.6% in Farmers Loop. Pre-thaw total nitrogen (% N) also varied across sites in the same manner, with the lowest being 0.11% in CRREL samples and highest at 2.15% in Farmers Loop samples. Combustible C and N data were not taken on post-thaw soils due to lack of sample. Gravimetric water content (GWC) varied between sites with the lowest being 61.6 % at CRREL and the highest at 313.8 % in Farmers Loop but thawing did not significantly change GWC across all sites.

**Table 2.** Pre-thaw and post-thaw permafrost abiotic soil properties. In the pre-thaw samples, “*” indicates that there were no subsample for the respective measurements (1 measurement per core due to sample limitation). Post-thaw samples were averaged across 4 replicates per core for pH, electrical conductivity (EC), and GWC measurements. The pH measurements were averaged for reporting with the following formula: −log_10[(ΣCi)/(n)] where C is the concentration of hydronium ion and n is the number of measurements (Boutilier and Shelton, 1980). % C and % N measurements were not taken on post-thaw soils due to sample limitation. Water was freely drained from thawing permafrost before post-thaw GWC measurements (see methods). Superscript letters after measurements indicate results from TukeyHSD tests using compact letter display for statistically significant differences.

| Pre-thaw permafrost |  |  |  |  |  |  |  | Post-thaw permafrost |  |  |  |
| --- | --- | --- | --- | --- | --- | --- | --- | --- | --- | --- | --- |
| Site | Core | pH* | EC<br>(mS/cm)* | %<br>C* | %<br>N* | %<br>C:N | %<br>GWC * | pH | EC<br>(mS/cm) | %<br>GWC | Change in % GWC<br>(pre-thaw to post-thaw) |
| CRREL | AT 3 | 4.84 | 350.9 | 3.6 | 0.18 | 20 | 61.6 | 4.83 ±<br>0.03<sup>b</sup> | 264.2 ±<br>57.7<sup>b</sup> | 16.8 ±<br>7.4 | - 44.8 ± 7.4 |
| CRREL | AT 4 | 4.91 | 245.3 | 1.8 | 0.11 | 16.4 | 91.9 | 4.88 ±<br>0.03<sup>b</sup> | 228.7 ±<br>91.7<sup>b</sup> | 6.6 ± 1.9 | - 85.3 ± 1.9 |
| FL | FL 1 | 4.71 | 441.6 | 37.9 | 1.9 | 19.9 | 313.8 | 4.71 ±<br>0.02<sup>b</sup> | 345.6 ±<br>37.5<sup>ab</sup> | 263.9 ±<br>23.1 | - 49.9 ± 23.1 |
| FL | FL 2 | 4.73 | 388.1 | 39.6 | 2.15 | 18.4 | 208.8 | 4.72 ±<br>0.05<sup>b</sup> | 431.7 ±<br>25.1<sup>ab</sup> | 146.6 ±<br>65.6 | - 62.2 ± 65.6 |
| Utqiagvik | BEO<br>1 | 4.41 | 206.3 | 30.9 | 1.67 | 18.5 | 176.5 | 4.06 ±<br>0.05<sup>a</sup> | 440.4 ±<br>84.5<sup>a</sup> | 150.8 ±<br>41.5 | - 27.5 ± 41.5 |
| Utqiagvik | BEO<br>2 | 4.16 | 328.6 | 21.8 | 1.07 | 20.4 | 116.9 | 3.94 ±<br>0.03<sup>a</sup> | 478.5 ±<br>104<sup>a</sup> | 129.3 ±<br>32.4 | + 12.4 ± 32.4 |

### 3.2 Pre-thaw and post-thaw bacterial abundance and cumulative respiration

Pre-thaw bacterial abundance was not significantly different across sites. However, the change in bacterial abundance with permafrost thaw was significantly different across sites (Figure 2) (Kruskal-Wallis p-value = 0.01). The CRREL permafrost tunnel had the lowest change in biomass following thaw and was significantly lower than Farmers Loop (Dunn adj. p-value = 0.03) and Utqiaġvik (Dunn adj. p-value = 0.02), which had the most change in biomass with thaw, likely due to variability between cores. Cumulative respiration was also significantly different between sites (Figure S1) (Kruskal-Wallis p-value = 0.0002). Cumulative respiration in CRREL tunnel samples were significantly lower than that of Farmers Loop (Dunn p-value = 0.0001) and Utqiaġvik (Dunn p-value = 0.007). We assessed the relationship between cumulative respiration and microbial abundance (gene copy number) (Figure 3). There was no significant correlation with cumulative respiration and pre-thaw copy number (Spearman’s rho = 0.11, p-value = 0.6). However, there was a significant positive correlation between cumulative respiration and post-thaw copy number (Spearman’s rho = 0.66, p-value = 0.001).

**Figure 2.**
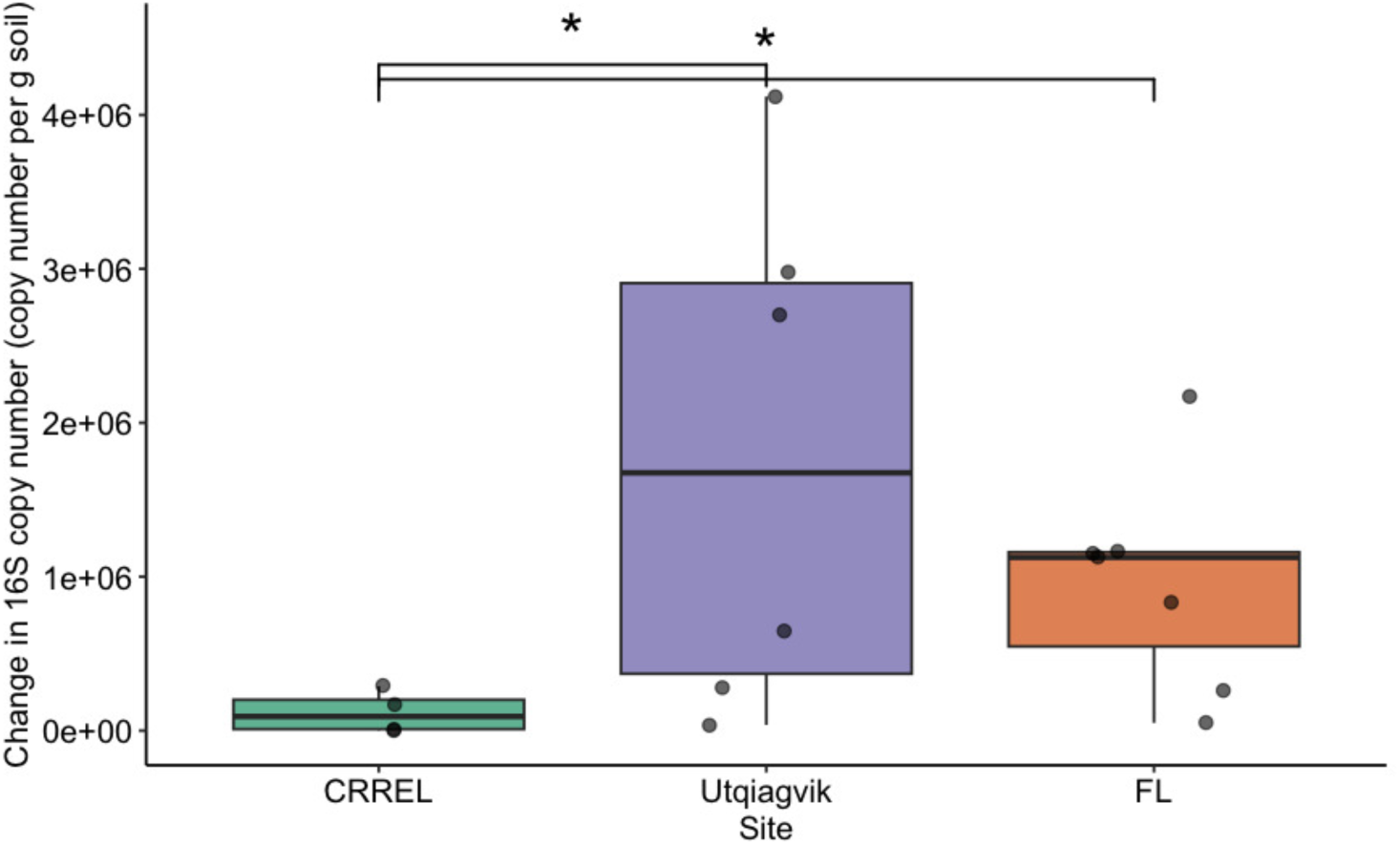
Bacterial abundance increased with thaw across all sites. Bacterial abundance was assessed via change in 16S copy number before and after thaw with significant differences between CRREL and Utqiaġvik (Dunn adj. p-value = 0.02) as well as CRREL and Farmers Loop (FL) (Dunn adj. p-value = 0.03). Boxplots depict the median value as a solid line and the upper and lower quartiles as the range of the box. Whiskers indicate the extent of the data and points represent raw data.

**Figure 3.**
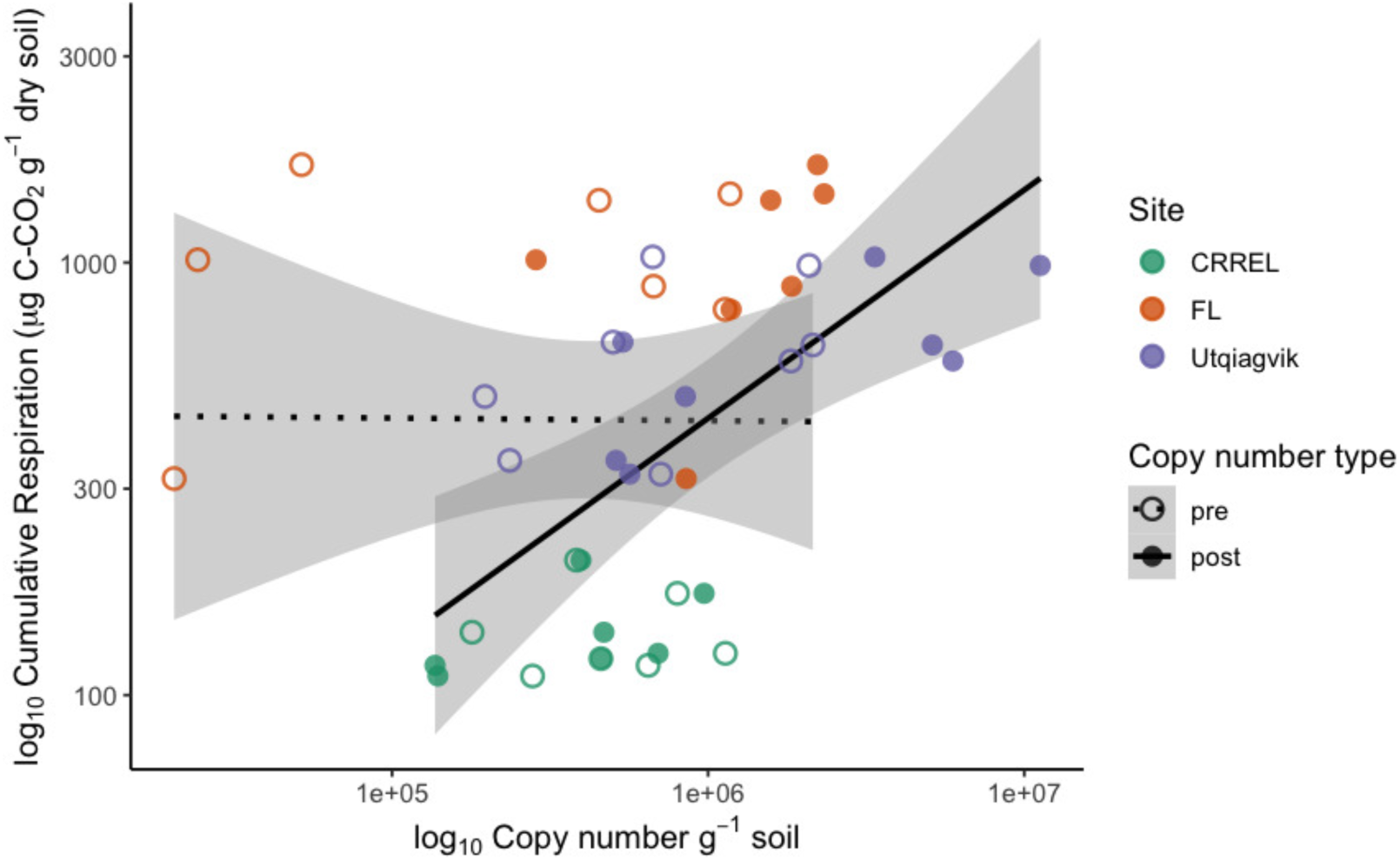
Cumulative respiration is positively correlated with post-thaw biomass. Pre-thaw Spearman correlation: rho = 0.11, *p* = 0.62; post-thaw Spearman correlation: rho = 0.66, *p* = 0.001.

### 3.3 Post-thaw microbiomes across three sites

Microbial communities retain site-specific composition before and after permafrost thaw (Figure 4). Permutational analysis of variance (PERMANOVA) showed significant effects of site (R^2^ =0.44, F = 18.9, p-value = 0.001) and thaw (R^2^ = 0.08, F =7.2, p-value = 0.001) on community composition. However, analysis of beta dispersion followed by a permutational test of multivariate homogeneity of groups (‘permutest’) indicated significant differences in dispersion among sites (F = 9.8, p-value = 0.001), suggesting that differences in community variability also contribute to observed site effects. Pairwise comparisons showed greater dispersion at Utqiaġvik compared to the CRREL tunnel and Farmers Loop (TukeyHSD adj. p-value = 0.001 for both).

**Figure 4.**
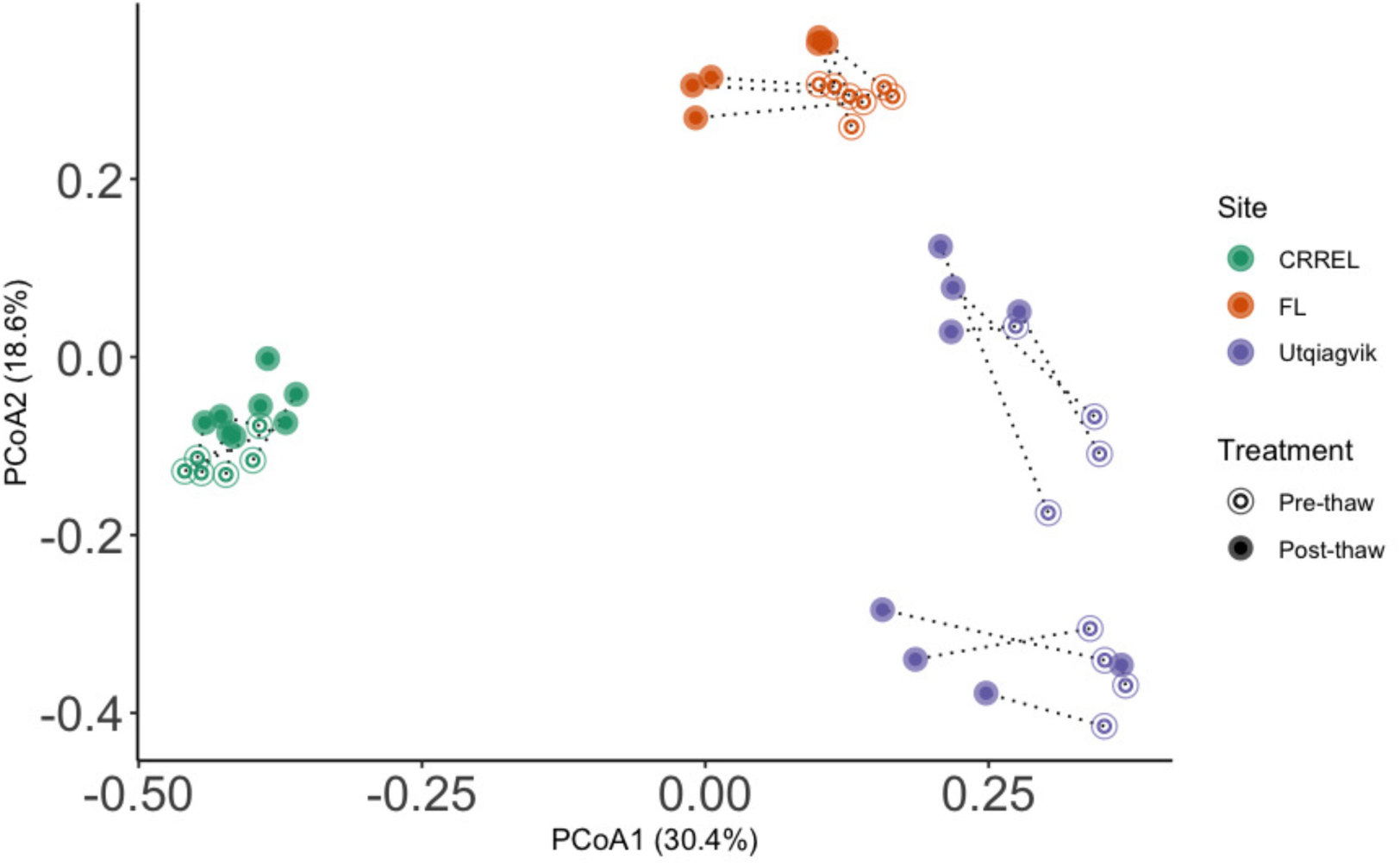
Microbial communities retain site-specific composition before and after permafrost thaw. PerMANOVA analysis revealed significant effects of site (R^2^ =0.44, F = 18.9, p-value = 0.001) and thaw (R^2^ = 0.08, F =7.2, p-value = 0.001) on microbial community composition.

Shannon diversity significantly differed between sites (Kruskal-Wallis p-value = 0.0001) and thaw treatment (Wilcoxon rank-sum test, W = 122, p = 0.007) such that diversity decreased with thaw (Figure 5a). Shannon values ranged from 3.25 to 5.25 in pre-thaw samples and from 0.67 to 4.89 in post-thaw samples. CRREL decreased by an average of 0.6 ± 0.25 post-thaw (ANOVA F = 5.171, p-value = 0.04), Farmers Loop decreased by an average of 0.7 ± 0.33 post-thaw (Kruskal-Wallis p-value not significant), and Utqiaġvik decreased by an average of 1.35 ± 0.25 post-thaw (Kruskal-Wallis p-value = 0.003). Similar trends to Shannon were observed with Simpson diversity as it was significantly different between sites (Kruskal-Wallis p-value = 0.0016) and thaw treatment (Wilcoxon rank-sum test, W = 81, p-value = 0.0001) (Figure 5b). Simpson values ranged from 0.83 to 0.99 in pre-thaw samples and from 0.24 to 0.97 in post-thaw samples. Further, Simpson diversity decreased with thaw as follows: CRREL decreased by an average of 0.032 ± 0.0085 (Kruskal-Wallis p-value = 0.002), Farmers Loop decreased by an average of 0.14 ± 0.048 post-thaw (Kruskal-Wallis p-value = 0.01), and Utqiaġvik decreased by an average of 0.23 ± 0.1 post-thaw (Kruskal-Wallis p-value = 0.006).

**Figure 5.**
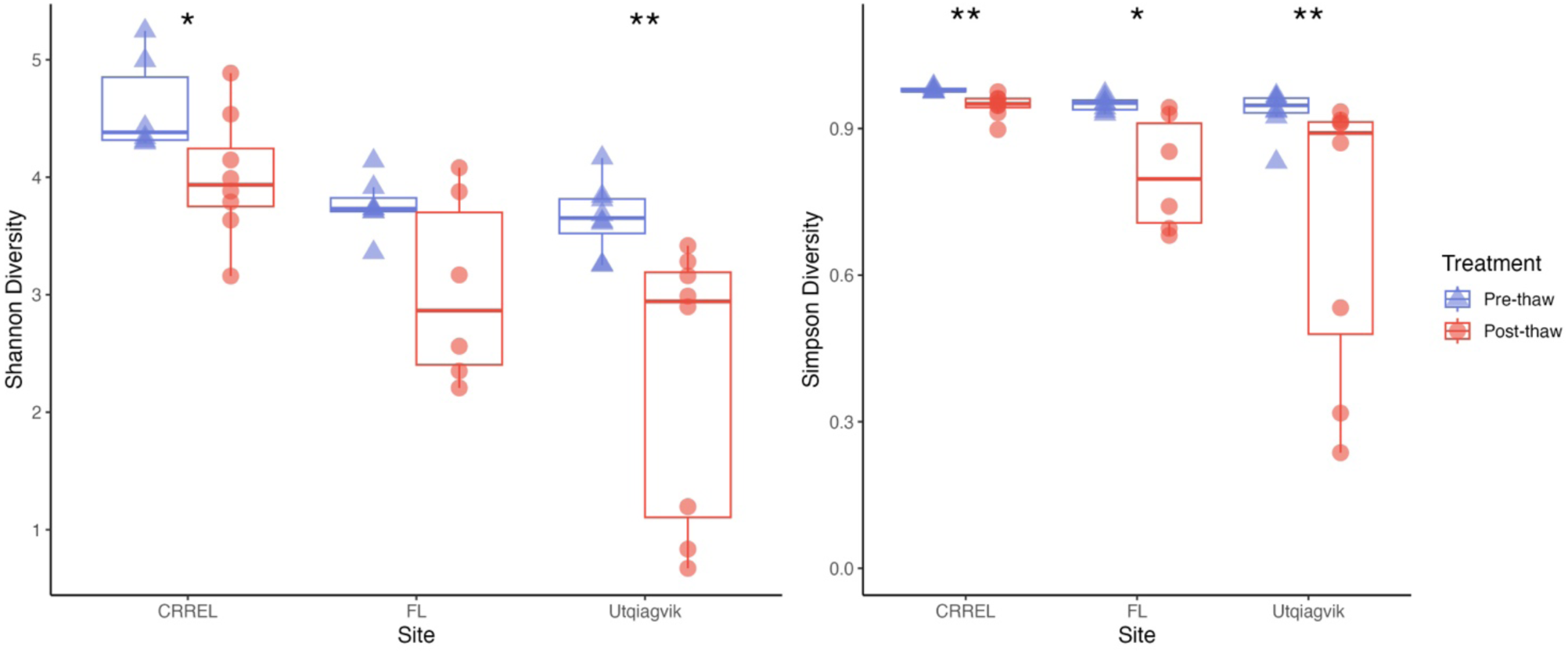
Alpha diversity decreases across all sites post-thaw. A) Shannon and B) Simpson diversity metrics for each site pre-thaw and post-thaw.

### 3.4 Analysis of thaw responders

Response ratio analysis was used to assess common thaw responders across all sites (Figure 6). A positive response ratio value indicates that a particular ASV responded positively to thaw— meaning at least a two-fold increase in abundance (Fan et al., 2022). We found a total of eight common thaw responders from all sites at the ASV level and identified them on the genus level: *Methylorosa*, *Burkholderia-Caballeronia-Paraburkholderia*, *Clostridium*, *Actimicrobium*, *Rugamonas*, *Massilia*, *Pseudomonas*, and Candidatus *Planktophila* (all p-values < 0.05). These responders spanned three phyla: Actinomycetota, Bacillota, and Pseudomonadota. We assessed positive thaw responders at each site individually and found 1153 ASVs (490 from the CRREL tunnel, 474 from Farmer’s Loop, and 189 from Utqiaġvik) from several other phyla increased in abundance following thaw (all p-values < 0.05) including common thaw-responder phyla. In CRREL tunnel samples, these included Acidobacteriota, Bacteroidota, Bdellovibrionota, Chloroflexota, Methanobacteriota, Planctomycetota, Thermodesulfobacteriota, and Verrucomicrobiota (Figure 7). In Farmer’s Loop samples, we observed increases in Acidobacteriota, Bacteroidota, Bdellovibrionota, Caldisericota, Gemmatimonadota, Myxococcota, Planctomycetota, Thermodesulfobacteriota, and Verrucomicrobiota (Figure 8). At Utqiaġvik, thaw was associated with increases in Acidobacteriota, Bacteroidota, Caldisericota, Deinococcota, Halobacteria, Methanobacteriota, Planctomycetota, Thermodesulfobacteriota, and Verrucomicrobiota (Figure 9).

**Figure 6.**
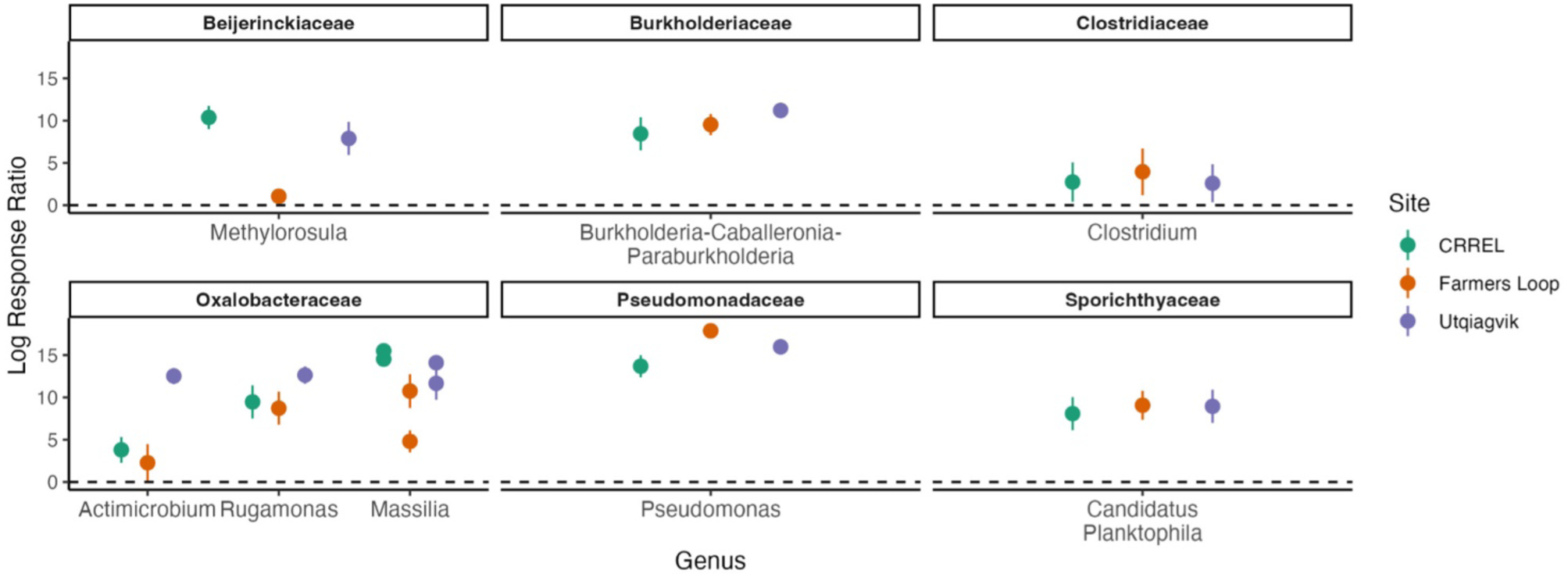
Positive thaw responder taxa, as determined by response ratio analysis, shared by all sites are from the phyla Actinomycetota, Bacillota, and Pseudomondadota (all p-values < 0.001). Family and genus are indicated on the figure titles and x-axis, respectively. Multiple points per genus indicate multiple species-level responses. Points represent the mean of replicates and error bars show the confidence interval.

**Figure 7.**
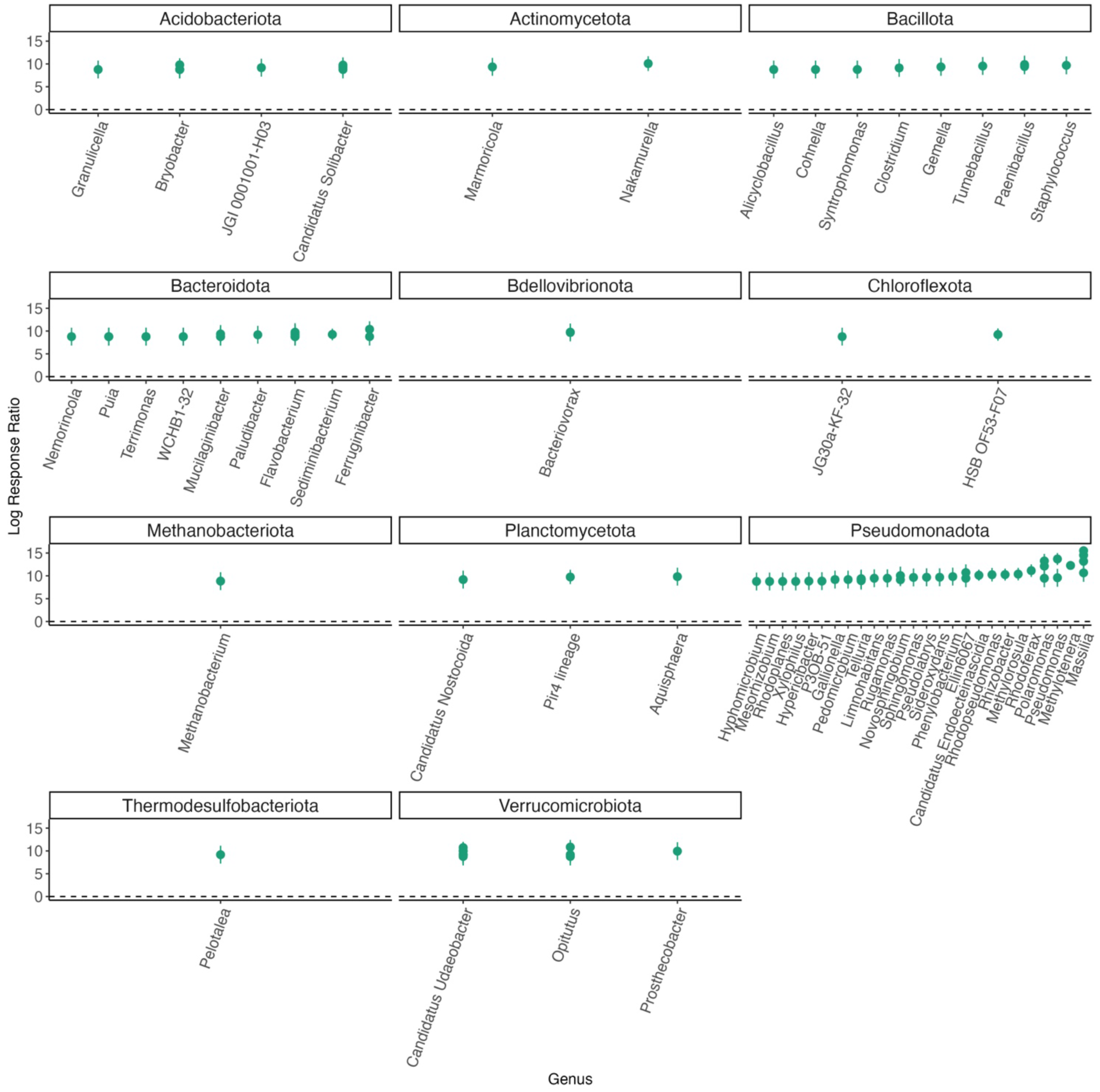
Positive thaw responder taxa, as determined by response ratio analysis, for the CRREL permafrost tunnel site. Phylum and genus are indicated in the figure titles and x-axis, respectively. Multiple points per genus indicate species-level responses. Points represent the mean of replicates and error bars show the confidence interval (all p-values < 0.001).

**Figure 8.**
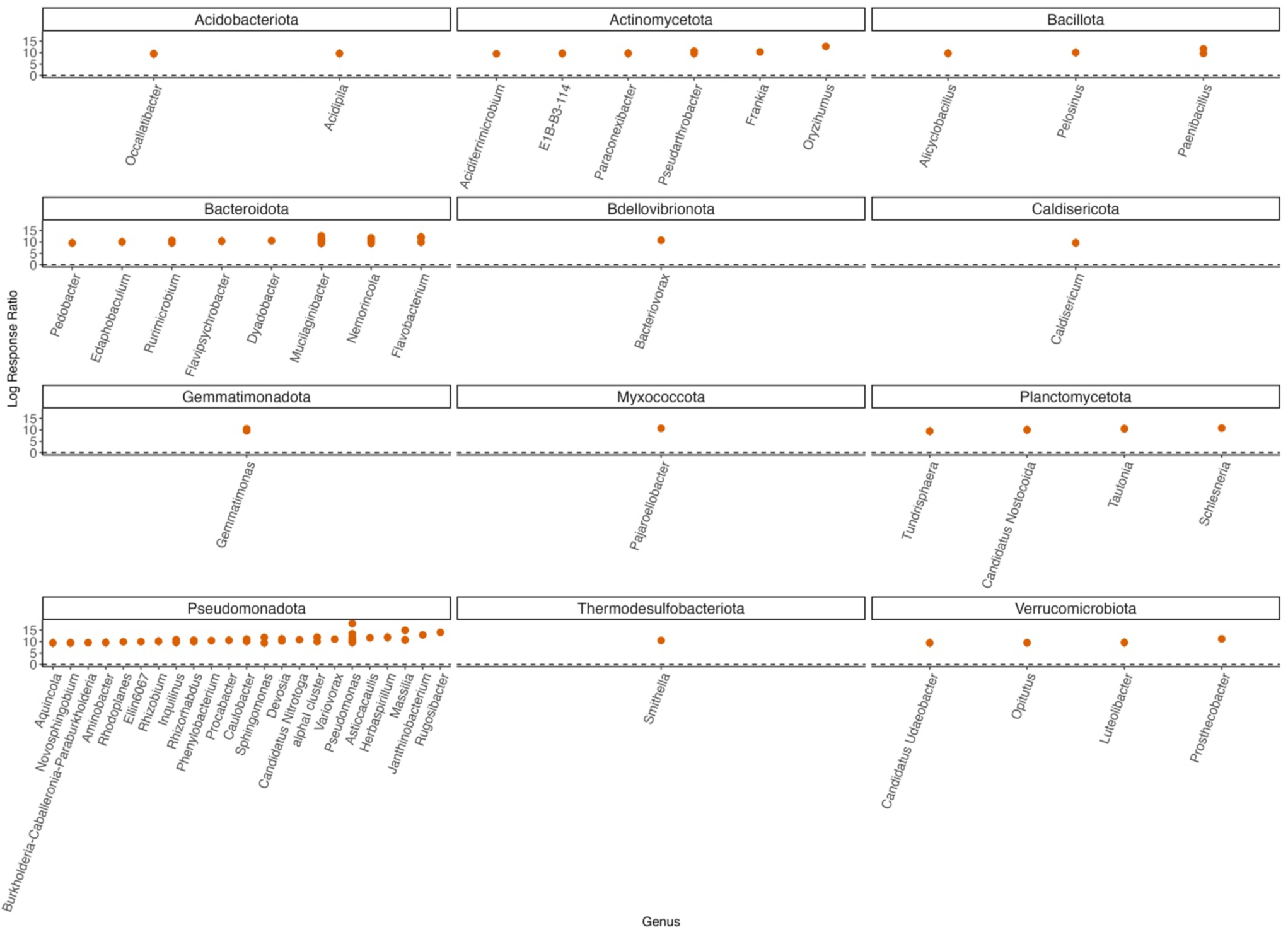
Positive thaw responder taxa, as determined by response ratio analysis, for the Farmer’s Loop site. Phylum and genus are indicated in the figure titles and x-axis, respectively. Multiple points per genus indicate species-level responses. Points represent the mean of replicates and error bars show the confidence interval (all p-values < 0.001).

**Figure 9.**
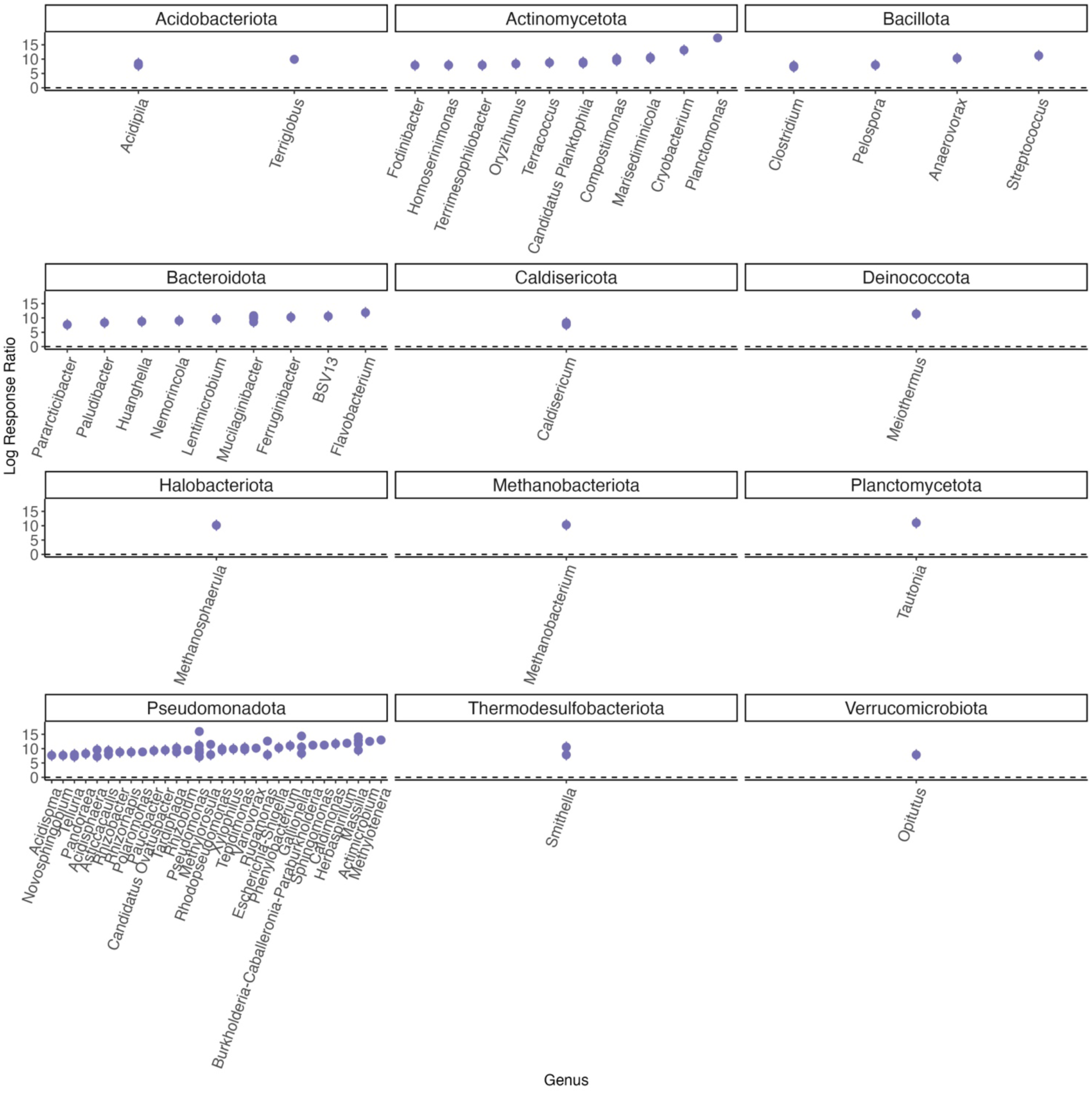
Positive thaw responder taxa, as determined by response ratio analysis, for the Utqiaġvik site. Phylum and genus are indicated in the figure titles and x-axis respectively. Multiple points per genus indicate species-level responses. Points represent the mean of replicates and error bars show the confidence interval (all p-values < 0.001).

## 4 Discussion

In this study, we investigated how permafrost thaw affects microbiome diversity, alters species abundance, and contributes to carbon flux across three distinct permafrost sites in Alaska through a microcosm incubation experiment. We found that (i) cumulative respiration and post-thaw microbial biomass were positively correlated, indicating that respiration has a strong relationship with new microbial growth; (ii) alpha diversity decreased following thaw across all sites; (iii) post-thaw microbiomes retain site-specific microbial community composition; (iv) certain phyla—Actinomycetota, Bacillota, and Pseudodesmonadota—consistently increased in abundance following thaw across all sites. These findings corroborate previous permafrost incubation studies and highlight key taxa that respond to the favorable conditions of thaw regardless of site.

### 4.1 Cumulative respiration is positively correlated with increase in bacterial abundance

Following thaw, bacterial abundance increased across all sites (Figure 2) likely due to the favorable conditions of thaw such as change in temperature and soil abiotic properties like % C, % N, pH, and EC. These abiotic factors largely influence the structure and function of soil microbial communities (Fierer, 2017; Horner-Devine et al., 2007) and have a direct effect on cumulative microbial respiration. In particular, permafrost thaw makes soil C more bioavailable (Coolen et al., 2011; Ernakovich et al., 2015; Waldrop et al., 2010) which is vulnerable to microbial decomposition (Ernakovich et al., 2017; Schuur et al., 2015; Waldrop et al., 2010; Zimov et al., 2006). It is worth mentioning that other microbes (fungi, archaea, etc.) are present in permafrost and contribute to the cumulative respiration in addition to bacteria. We found that the increase in bacterial abundance was positively correlated with cumulative respiration (Figure 3) indicating that microbial growth and the larger biomass pool is driving permafrost C losses. Carbon content in permafrost is one of the key drivers of microbial respiration (Dutta et al., 2006; Schädel et al., 2016; Zimov et al., 2006) and because of that, as permafrost thaws, there are higher rates of microbial activity and increased respiration rates (Ernakovich et al., 2017; Mackelprang et al., 2011; Monteux et al., 2020; Schädel et al., 2016; Schuur et al., 2015). Other studies have indicated that the amount of organic substrate in permafrost has the potential to dictate microbial abundance and respiration (Müller et al., 2018; Stackhouse et al., 2015). Indeed, the amount of organic matter in the soil appears to be the dominant factor influencing bacterial abundance and respiration responses to thaw between our sites, as CRREL samples contained less C (∼ 3%) and N (∼ 0.15%) (on a per mass basis) than Farmer’s Loop (%C ∼38%, %N ∼ 2%) and Utqiaġvik (%C ∼ 26%, %N ∼ 1.5) for pre-thaw conditions (Figure S2, S3). Taken together, our findings of the positive relationship between microbial abundance and cumulative respiration supports other findings that the increase in soil microbial activity with thaw is due to the growth of the microbial biomass (Coolen & Orsi, 2015; Ernakovich et al., 2017; Schädel et al., 2016; Schuur et al., 2015).

### 4.2 Post-thaw microbiomes are site-specific but contain common taxa with a consistent response to thaw

Following thaw, the post-thaw microbiomes across our three sites decreased in diversity, changed in composition, and increased in absolute abundance, all of which have the potential to alter community function. This is consistent with previous studies that found that microbiome composition, functional gene abundance, and metabolic pathways change dramatically with permafrost thaw (Coolen & Orsi, 2015; Doherty et al., 2020; Mackelprang et al., 2011; Tuorto et al., 2014; Waldrop et al., 2025). Our findings that alpha diversity decreased with thaw (Figure 5) is consistent with other studies (Thurston et al., 2025). Farmers Loop and Utqiaġvik had similar microbial diversity, as assessed with the Shannon and Simpson metrics. This may be due to similar soil properties (e.g., soil C and N). However, microbial diversity was significantly different between CRREL and both Farmers Loop and Utqiaġvik, which may lend further support to the relationship between diversity and soil C and N. Further, this decrease in diversity is likely driven by abiotic changes associated with thaw, particularly shifts in soil pH. Previous work has shown that Shannon diversity is unimodally related to soil pH, with lower diversity in more acidic conditions (Waldrop et al., 2023). Consistent with this, we found that pH generally declined across all samples post-thaw. While diversity decreased, microbial biomass increased indicating that there are fewer, more abundant taxa which is common in disturbance events (Ernakovich et al., 2022; Mackelprang et al., 2011; Seitz et al., 2021). Further, the combination of responders and lower diversity post thaw makes a strong case for certain taxa driving biogeochemistry and greenhouse gas production.

Understanding common microbial responses to thaw is imperative for predicting the structure and function of post-thaw microbial communities. The power of our cross-site study is that, despite site specific differences in the pre- and post-thaw community compositions, we found common thaw responders (analyzed at the ASV level and identified at the genus level) that increased at least two-fold in abundance following thaw. We define thaw responders as bacterial taxa that increase in abundance post-thaw. An increase post-thaw is likely due to the favorable conditions that permafrost thaw presents (increase in labile carbon, moisture, etc.). There were a number of taxa from the phyla Actinomycetota, Bacillota, and Pseudomonadota that were common thaw responders across sites (Figure 6). The phyla of thaw responder results observed in this study are consistent with other studies Actinomycetota: (Chen et al., 2021; Mackelprang et al., 2011; Monteux et al., 2020; Scheel et al., 2023; Stackhouse et al., 2015), Bacillota (formerly known as Firmicutes): (Barbato et al., 2022; Chen et al., 2021; Coolen & Orsi, 2015; Monteux et al., 2020; Stackhouse et al., 2015; Yang et al., 2021), Pseudomonadota (D. Kim et al., 2025). These responders belong to the genera: *Methylorosula*, *Burkholderia-Caballeronia-Paraburkholderia*, *Clostridium*, *Actimicrobium*, *Rugamonas*, *Massilia*, *Pseudomonas*, and *Candidatus Planktophila*. Additionally, each site also had its own set of taxa that responded to thaw (Figures 7, 8, 9). Interestingly, the site-specific thaw responder results align with recent findings of the same phyla present in the atmosphere as bioaerosols above major permafrost thaw zones. These airborne microbes actively contribute to the Arctic aerobiome which can lead to cloud formation, alter precipitation patterns, and affect radiative forcing (Nieto-Caballero et al., 2025). Here, we focus on the roles and characteristics of each of these common thaw responders, and what their response may mean for post-thaw permafrost microbiome composition under aerobic conditions. These responders vary in their functional roles and characteristics, as some of these genera are known for their pigment producing properties, pathogenic potential, promotion of plant growth, and contributions to biogeochemical cycling.

#### 4.2.1 Actinomycetota

Candidatus *Planktophila*—freshwater bacterioplankton—increased in all sites following thaw. This genus has no known isolates to date, but has been studied in a stable mixed culture (Jezbera et al., 2009) and described as amino acid prototrophs (Neuenschwander et al., 2018). It is likely that this genus increases following thaw due to increased water flow, which can promote the formation of thaw lakes where this genus is largely abundant (Bomberg et al., 2019; Vigneron et al., 2019). Indeed all of our permafrost samples were high in ice content and can contribute to large thermokarst features with thaw.

#### 4.2.2 Bacillota

*Clostridium*, a genus of the phylum Bacillota is gram-positive, spore-forming, anaerobic bacteria known for the potential pathogenicity of some of its species. It is likely that *Clostridium* were dormant under the frozen conditions in permafrost, as they can enter dormancy through sporulation when conditions are unfavorable*. Clostridium* has been previously found to increase in abundance with thaw (Barbato et al., 2022; Chen et al., 2021; Coolen & Orsi, 2015; Yang et al., 2021), while other studies have noted their decrease (Kim et al., 2025). It is likely that the increase with permafrost thaw is due to germination of spores, as they respond to favorable conditions for microbial growth and activity. This genus also includes several pathogenic strains that infect humans and animals and may pose a threat to ecosystems during and after thaw.

#### 4.2.3 Pseudomonadota

*Pseudomonas* is a genus of gram-negative, aerobic bacteria found in a wide variety of environments such as soil, water, plant seeds, and animal hosts (Palleroni, 2008). A number of *Pseudomonas* strains have pathogenic potential. Recent work has revived potato plant pathogens from thawing permafrost, as these previously dormant bacteria increased with thaw (Kim et al., 2025).

For instance, *Methylorosula* is a gram negative, aerobic, motile bacteria within the family *Beijerinckiaceae*. It has one known species *Methylorosula polaris*, originally isolated from acidic tundra wetland soils of Northern Russia. This genus is facultatively methylotrophic and not capable of nitrogen fixation (Berestovskaya et al., 2012). Instead, *Methylorosula* are known for their metabolic versatility because they can synthesize most carbon compounds such as sugars, polysaccharides, ethanol, and amino acids, in addition to one-carbon compounds like methanol and methylamines (Berestovskaya et al., 2012). It is likely that this genus is a thaw responder because these compounds are degraded during permafrost thaw via decomposition done by other microbes. *Methylorosula’s* ability to process single carbon compounds may aid in reducing methane emissions from permafrost, as they may limit substrates available to methanogens and regulate carbon fluxes.

*Burkholderia-Caballeronia-Paraburkholderia* (BCP) is one of the common thaw responders and has been previously noted to respond positively to thaw (Scheel et al., 2023). BCP refers to three closely related genera within the *Burkholderiaceae* family. The genus *Burkholderia* was recently subdivided into three clades: *Burkholderia* (clade I) which consists mostly of plant and animal pathogens, *Caballeronia* (clade II) which consists of environmental isolates, and *Parkaburkholderia* (clade III) to distinguish plant growth promoting and root-associated bacteria (Madhaiyan et al., 2021; Sawana et al., 2014). BCP are gram-negative bacteria and are known for their plant growth promoting properties as a rhizobacteria (Madhaiyan et al., 2021). BCP contribute to organic matter decomposition releasing CO_2_ and some contribute to nitrogen cycling (*Paraburkholderia*). Some *Paraburkholderia* are capable of nitrogen fixation and shown to be able to suppress phytopathogens which may benefit plants from pathogens in thawing permafrost ecosystems. The contribution of nitrogen fixation and plant growth promotion could enhance vegetation with permafrost thaw, which could end up storing more carbon in the ground.

*Actimicrobium* is a gram-negative, strictly aerobic non-motile genus of the family *Oxalobacteriacee*. It has one known isolate *Actimicrobium antarcticum*, originally isolated from Antarctic coastal seawater (Kim et al., 2011) and was recently found to be in high abundance in Antarctic glacier boreholes (Cui et al., 2025). Like *Actimicrobium*, *Rugamonas* was also originally isolated from Antarctica, but from freshwater samples. *Rugamonas* is described as a genus of gram-negative aerobic bacteria. Recent work has shown that members of this genus produce violacein, a microbial antibiotic pigment (Choi et al., 2015; Sedláček et al., 2021). Similarly, *Massilia*, another positive thaw responder also produces violacein (Sedláček et al., 2021).

## 5 Conclusions

As permafrost thaws, the microbiome composition, abundance, and diversity changes. Collectively, these changes have the potential to accelerate climate change. Our results identify common positive thaw responders that are likely contributing to the increase in respiration with thaw. Additionally, these particular responders are known for their antimicrobial pigments, pathogenic potential, and plant growth promoting properties indicating that an increase in these taxa can alter community dynamics, lead to infectious outbreaks, and promote vegetation, all of which have implications for the carbon cycle. Future research is needed to further elucidate the function of these positive thaw responders, the post-thaw microbiome, and in particular—its pathogenic potential all of which can inform predictions for ecosystem change due to permafrost thaw.

## Author Contributions

J.M.O: Writing – original draft, review and editing. N.D.B: Writing – review and editing, Investigation. H.H.M: Writing – review and editing, Methodology, Supervision, Formal Analysis, Validation. K.L.S: Data curation, Writing – review and editing, T.A.D: Writing – review and editing, Resources. R.A.B: Writing – review and editing, Conceptualization, Methodology, Funding acquisition, Resources. J.G.E: Writing – review and editing, Conceptualization, Methodology, Supervision, Project administration, Validation, Funding acquisition, Resources, Visualization.

## Acknowledgments

We would like to give a special thanks to Dr. Vanessa Bailey, Julie Bobyock, Lukas Bernhardt, Dr. Taylor Sullivan, Dr. Andy Parsekian, and Dr. David Olefeldt for their support as well as past and present members of the Ernakovich Lab and Center for Soil Biogeochemistry and Microbial Ecology (Soil BioME) at the University of New Hampshire. We would also like to thank Dr. Ted Schuur and Dr. Christina Schädel for organizing the Permafrost Carbon Network (NSF PLR Grant# 1931333).

## Data Availability Statement

The datasets presented in this study can be found in online repositories. The names of the repository/repositories and accession number(s) can be found below: https://www.ncbi.nlm.nih.gov/sra/PRJNA1284569

