## Supplementary Information for "Consistent microorganisms respond during aerobic thaw of Alaskan permafrost soils"

Supplementary Material

### **1 Supplementary Figures**

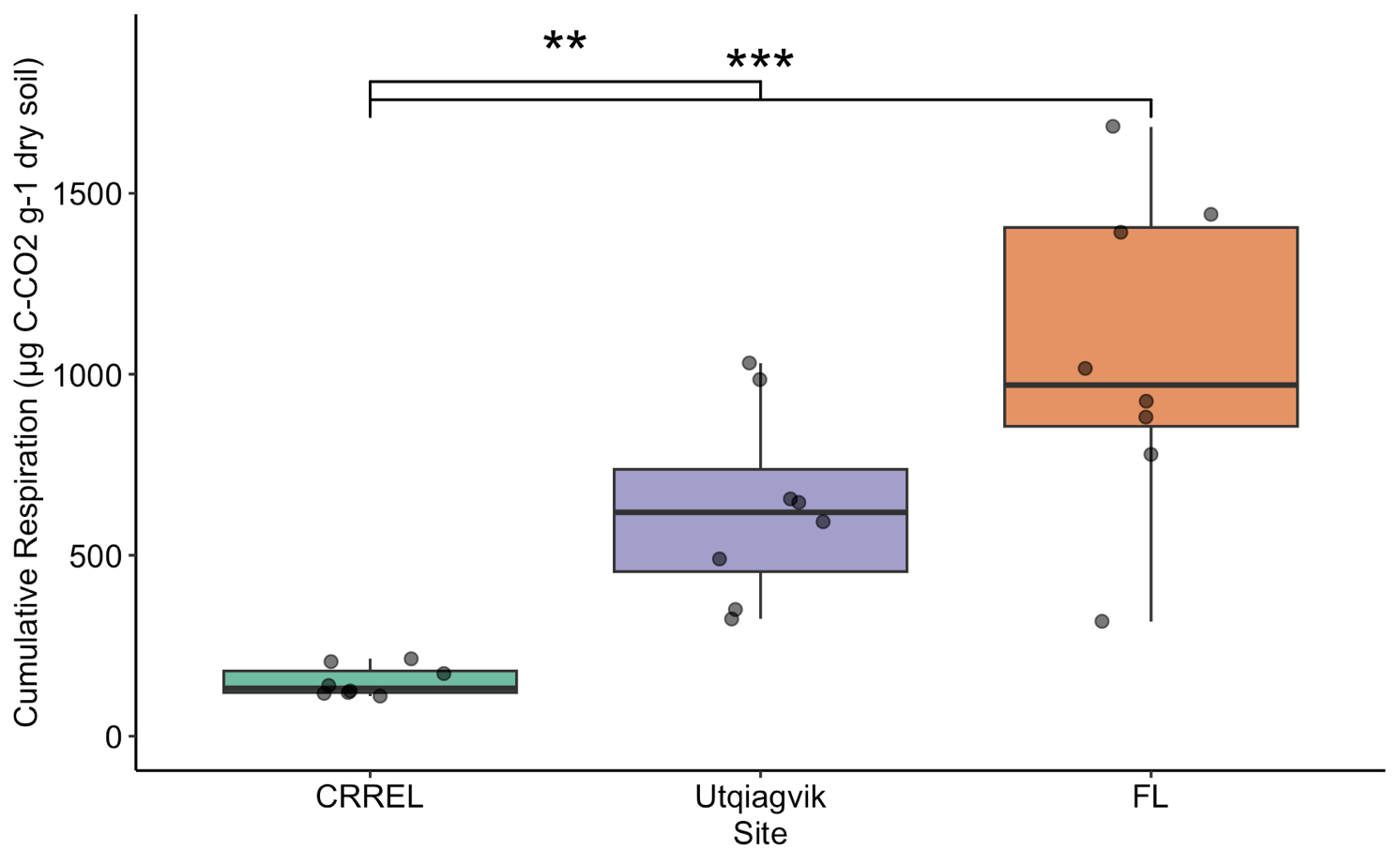

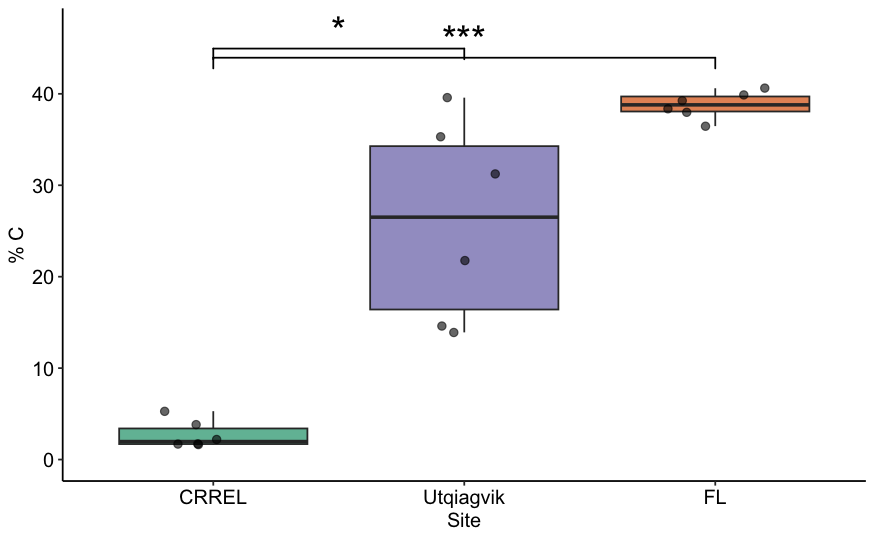
**Supplementary Figure 1.** Cumulative respiration during thaw varies among sites (Kruskal-Wallis p-value = 0.0002) with significant differences between CRREL and Farmer’s Loop (FL) (Dunn p-value = 0.0001) as well as CRREL and Utqiagvik (Dunn p-value = 0.007). Boxplots depict the median value as a solid line and the upper and lower quartiles as the range of the box. Whiskers indicate the extent of the data and points represent raw data.

**Supplementary Figure 2.** Total carbon varies by site. Significant differences between CRREL and Utqiagvik (Dunn p-value = 0.03) and CRREL and Farmer’s Loop (FL) (Dunn p-value = 0.0002). Boxplots depict the median value as a solid line and the upper and lower quartiles as the range of the box. Whiskers indicate the extent of the data and points represent raw data.

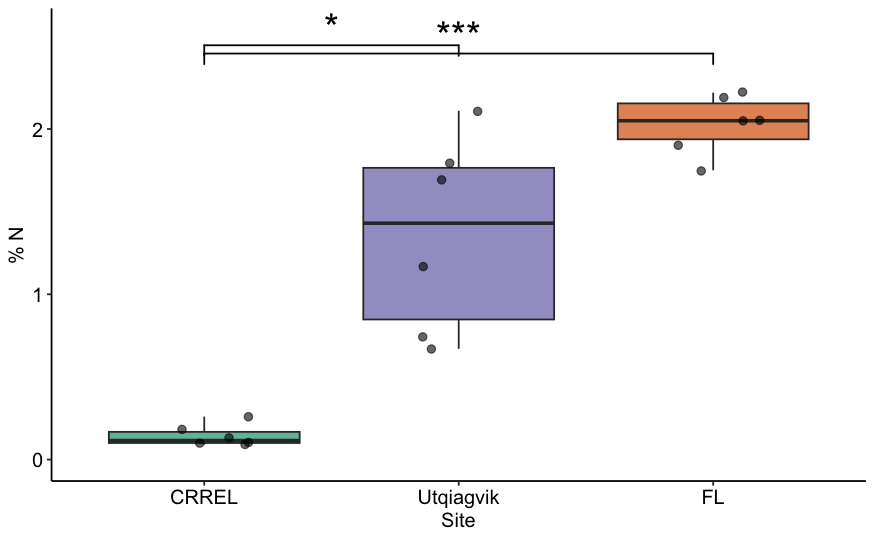

**Supplementary Figure 3**. Total nitrogen varies by site. Significant differences between CRREL and Utqiagvik (Dunn p-value = 0.02) and CRREL and Farmer’s Loop (FL) (Dunn p-value = 0.0002). Boxplots depict the median value as a solid line and the upper and lower quartiles as the range of the box. Whiskers indicate the extent of the data and points represent raw data.
